# REPIC: A database for exploring *N*^6^-methyladenosine methylome

**DOI:** 10.1101/2019.12.11.873299

**Authors:** Shun Liu, Chuan He, Mengjie Chen

## Abstract

The REPIC (<u>R</u>NA <u>Epi</u>transcriptome <u>C</u>ollection) database records about 10 million peaks called from publicly available m^6^A-seq and MeRIP-seq data using our unified pipeline. These data were collected from 672 samples of 49 studies, covering 61 cell lines or tissues in 11 organisms. REPIC allows users to query *N*^6^-methyladenosine (m^6^A) modification sites by specific cell lines or tissue types. In addition, it integrates m^6^A/MeRIP-seq data with 1,418 histone ChIP-seq and 118 DNase-seq data tracks from the ENCODE project in a modern genome browser to present a comprehensive atlas of m^6^A, histone modification sites and chromatin accessibility regions. REPIC is accessible at http://epicmod.uchicago.edu/repic.

## Background

Over a hundred fifty types of chemical modifications have been identified in messenger RNAs (mRNAs) and non-coding RNAs (ncRNAs)[1]. Among them, *N*^6^-methyladenosine (m^6^A) is characterized as the most abundant and reversible RNA internal modification[2, 3]. Increasing studies suggest that m^6^A has emerged as a critical regulator of post-transcriptional gene expression programs and involves with many cellular activities including splicing[4], translation efficiency[5], stability[6], export and cytoplasmic localization[7] of the modified mRNAs. Further, m^6^A also impacts a series of physiological processes related to proliferation[8], development[9], neurogenesis[10], circadian rhythm[11] and embryonic stem cell differentiation[12].

With the development of next-generation sequencing (NGS) technologies, several high-throughput sequencing strategies (m^6^A-seq or MeRIP-seq[13, 14], PA-m^6^A-seq[15], m^6^A-LAIC-seq[16], miCLIP[17, 18], m^6^A-REF-seq[19], MAZTER-seq[20] and DART-seq[21]) have developed to explore distributions and quantitative features of m^6^A modifications across the entire transcriptome, paving the path for understanding their biological functions. These methods, especially m^6^A/MeRIP-seq, have been largely adopted to profile the m^6^A marks in a variety of cell lines and tissue types from multiple species. To better explore m^6^A datasets with increasing complexity, several databases (RMBase v2.0[22], MeT-DB v2.0[23], CVm6A[24]) and webservers (RNAmod[25], WHISTLE[26], SRAMP[27]) have been developed to organize and integrate existing resources. Among these, RMBase v2.0 integrated published m^6^A peaks from more than five RNA modifications, RBP binding sites, and single nucleotide polymorphisms, while MeT-DB v2.0 and CVm6A focused on m^6^A peaks called by applying their own pipelines to raw m^6^A sequencing data. However, these databases have limitations. It has been shown that distinct m^6^A patterns occur in different developmental stages or tissue types, implying their dynamic regulation in a tissue-dependent manner[28]. Unfortunately, all the above databases except CVm6A simply combined m^6^A peaks across datasets without considering the cell type- or tissue-specificity. Furthermore, recent studies uncovered the associations of m^6^A modifications with promoters[29–31] and histone marks[32, 33], offering new insights into potential regulation pathways and underlying mechanisms, through which m^6^A could influence transcriptional regulation and gene expression. However, to our knowledge, m^6^A modifications and epigenomic data have not been curated well together. New bioinformatic tools are needed for processing, analyzing and visualizing such data integration.

Here, we present the REPIC (<u>R</u>NA <u>Epi</u>transcriptome <u>C</u>ollection) database, which currently focuses on integrating m^6^A modifications with the ENCODE epigenomic data. The m^6^A modification peaks are generated by re-processing publicly available m^6^A-seq and MeRIP-seq datasets using a unified customized pipeline. REPIC allows users to query m^6^A modification sites by cell lines or tissue types with a user-friendly interface and provides a built-in genome browser for visualization. Overall, REPIC is designed as a new resource to explore cell/tissue-specific m^6^A modifications and investigate potential interactions between m^6^A modifications and histone marks or chromatin accessibility.

## Construction and content

The REPIC database collected m^6^A modifications and epigenomic sequencing data from different species. We designed a modern and user-friendly web portal for querying m^6^A modification sites and an interactive genome browser empowered by GIVE[34] for data visualization (**Fig. 1a**). The web application of the REPIC database was constructed using Apache v2.4.18, MySQL v5.7.25 and PHP v7.2.14. The source of raw data and data processing procedure were shown in **Fig. 1b**. To better disseminate the resource and facilitate downstream analysis, we provide curated data that can be downloaded from the database website.

**Fig. 1.**
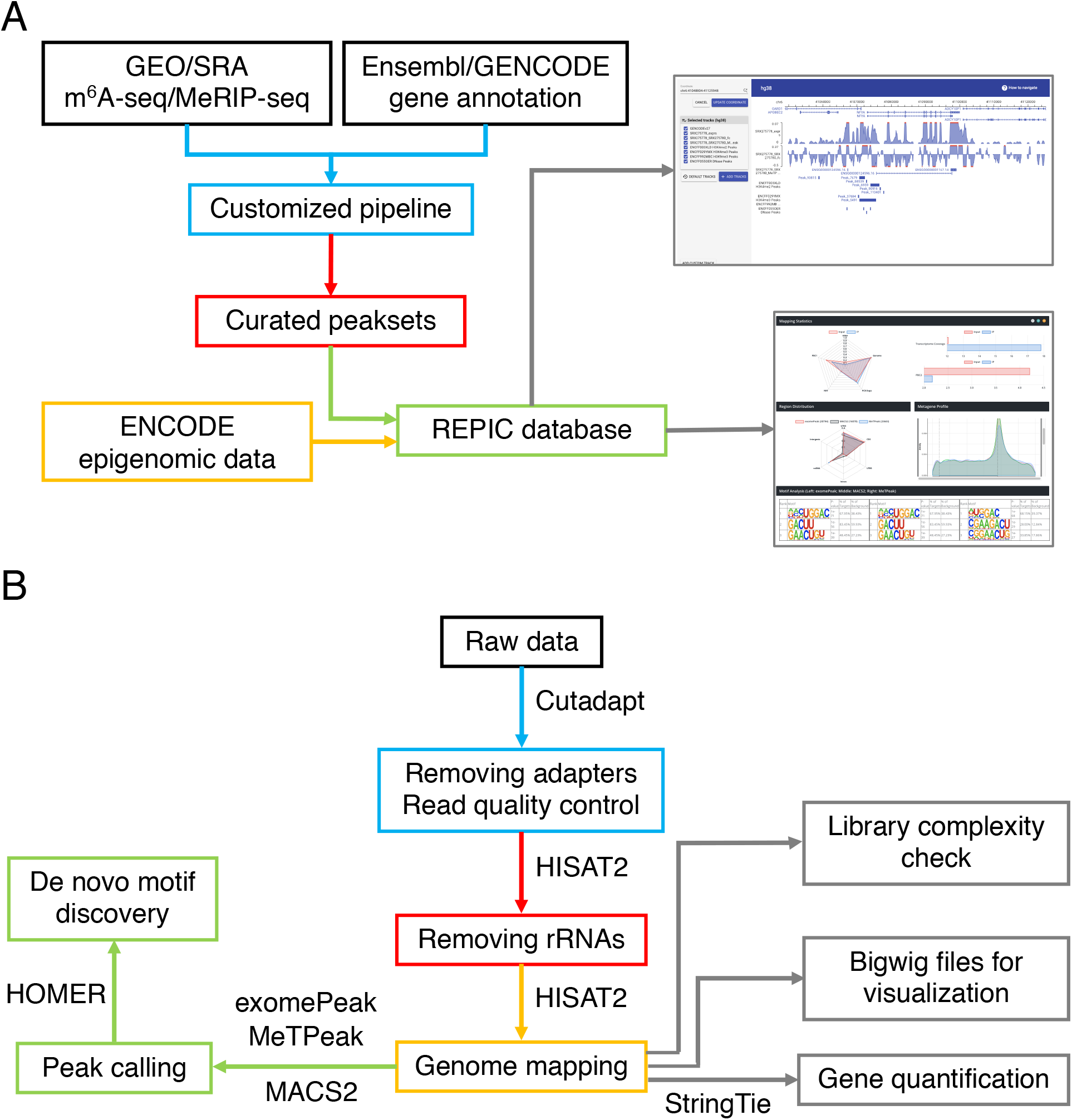
a) Overall design of the REPIC database. b) Schema of the customized pipeline for m^6^A-seq or MeRIP-seq data processing.

### High-throughput sequencing data

Raw m^6^A-seq and MeRIP-seq data were manually collected through literature searching and then retrieved from the Gene Expression Omnibus (GEO) and Sequence Read Archive (SRA). In total, 607 m^6^A-seq and 544 meRIP-seq run data were obtained from SRA. After merging different runs in the same experiment and excluding unpaired input-IP samples, 672 samples which consisted of 339 paired input-IP data from 49 studies, covering 61 cell lines or tissue types in 11 organisms, were used for the database construction (**Table S1**). For epigenomic data, a total of 118 DNase-seq peak sets from 29 cell lines or tissue types, and 1,418 histone ChIP-seq peak sets from 27 histone marks in 22 cell lines or tissue types in human and mouse, matching with curated m^6^A modification data, were downloaded from the ENCODE website (**Table S2 and S3**).

### Genome annotation data

The genome sequences and gene annotations of human and mouse were acquired from UCSC Genome Browser[35] and GENCODE[36], respectively. *Arabidopsis thaliana* genome sequences and gene annotations were obtained from the Arabidopsis Information Resource (TAIR)[37]. Others were downloaded from the Ensembl website[38]. The widespread-used versions of genome sequences and gene annotations of all 11 organisms were chosen for further analysis (**Table S4**).

### Raw m^6^A-seq and MeRIP-seq data reprocessing

The aforementioned 339 paired input-IP data were re-processed by our customized pipeline (https://github.com/shunliubio/easym6A) (**Fig. 1b**). Briefly, adapters of raw sequencing data were clipped by Cutadapt v1.15[39]. Reads longer than 15 nt after trimming were first mapped to ribosomal RNAs (rRNAs) by HISAT2 v2.1.0[40]. All unmapped reads were then aligned to genomes using HISAT2 v2.1.0 with default parameters. To check library complexity, PCR duplicates were evaluated by MarkDuplicates from Picard v2.17.10[41]. Then PCR Duplicate Proportion (PDP) was defined as the number of PCR duplicates divided by total number of mapped reads. Another three indexes, Non-Redundant Fraction (NRF), PCR Bottlenecking Coefficients 1 (PBC1) and 2 (PBC2), were measured according to the ENCODE standards. Input samples from m^6^A-seq and MeRIP-seq data were used to estimate gene expression levels by StringTie v1.3.4d[42]. If the library type is strand-specific, we further divided the sequence alignment data by strands. Log2 fold enrichment of m^6^A was calculated and gene expression levels were reported in Bins Per Million mapped reads (BPM) using deepTools v3.0.2[43]. exomePeak[44], MeTPeak[45] and MACS2 v2.1.1[46] were used to detect peaks. Finally, HOMER v4.9[47] was used for motif enrichment analysis based on top 2,000 peaks ranked by their fold enrichment levels.

## Utility and discussion

### Quality evaluation of m^6^A-seq and MeRIP-seq data

We applied our pipeline to re-process all collected m^6^A-seq and MeRIP-seq samples. As rRNAs provide little information about transcriptome, we first interrogated the rRNA content in each sample. As a result, rRNAs were less than 30% of total reads in 566 samples (85.0% of the total), while 371 samples (55.7% of the total) contained rRNA reads below 5% (**Fig. 2a**), suggesting that most of samples were not subjected to rRNA contamination. Next, we examined the counts of reads mapped to the genomes after filtering out rRNA reads. As a result, 571 (85.7%) were shown of high quality with a genome mapping ratio greater than 75% (**Fig. 2b**). 16 human and 22 mouse samples with a low genome mapping ratio (< 60%) were detected as virus infection, vector or mycoplasma contamination or other unknown conditions.

**Fig. 2.**
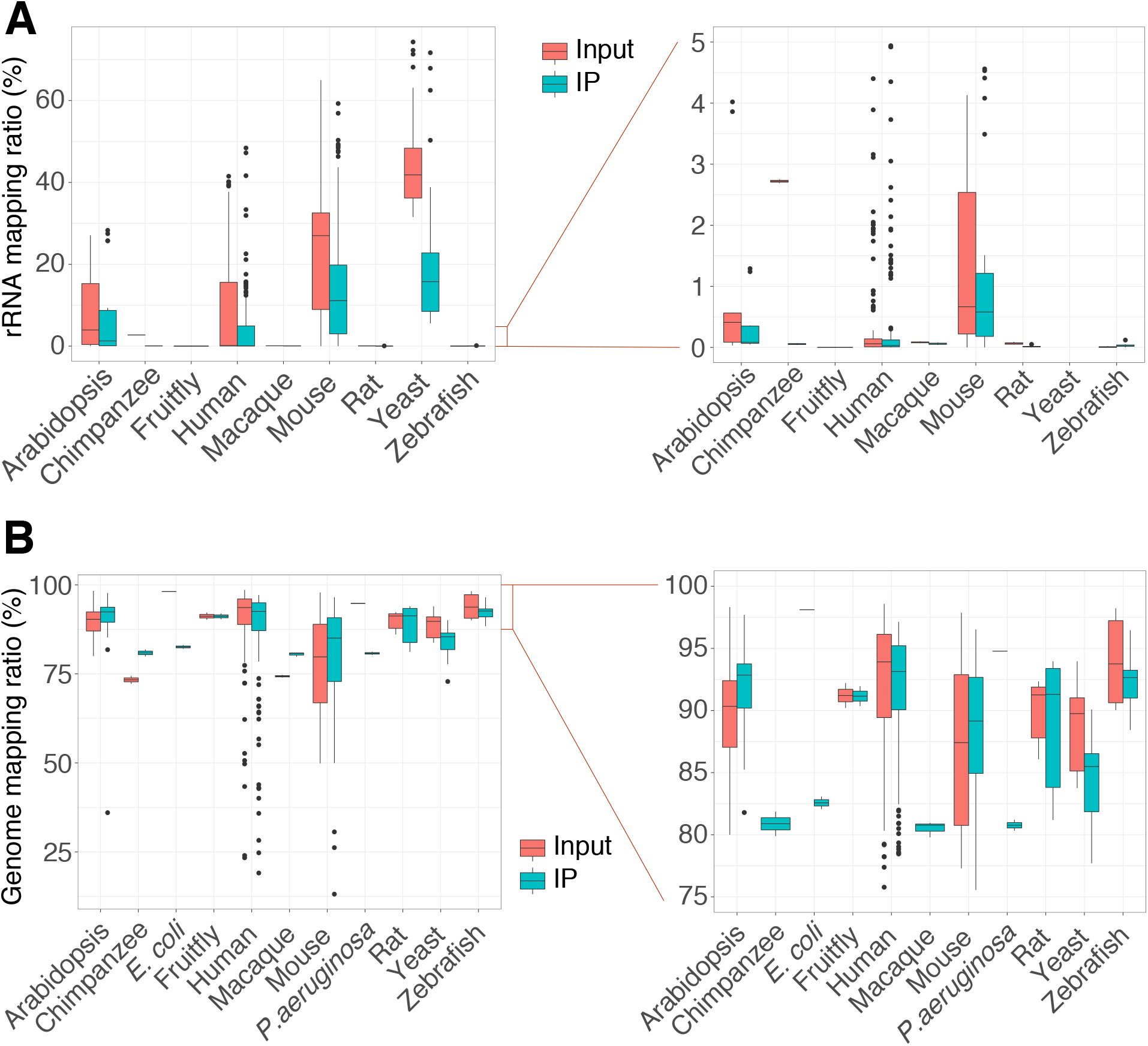
The quality of m^6^A-seq or MeRIP-seq reads mapping. Boxplots depicting the distribution of input and IP samples within rRNA (a) and genome (b) mapping ratios. Both the left panels showing the whole range of the ratios and the right panels of (a) and (b) zooming in the range from 0-5% and 75-100%, respectively.

To further evaluate data quality, we assessed the library complexity of all samples by four metrics, PDP, NRF, PBC1 and PBC2, with the latter three defined by the ENCODE project[48]. PDP indicated that about 75% of the samples contained PCR duplicates over 50% (**Fig. S1a**), while NRF showed that about 25% of the samples had distinct uniquely mapping reads more than 50% (**Fig. S1b**). Both PDP and NRF implied that multiple reads in the same positions of the genomes were prevalent in the samples. However, whether to remove them as PCR duplicates is an open question since it is difficult to distinguish between PCR amplification artifacts and real transcriptional events using current computational methods. Direct removal of duplicated reads with the same mapping coordinates may introduce unwanted bias[49, 50]. Therefore, our pipeline kept duplicated reads for downstream analysis. Unlike PDP and NRF, about 90% of the input samples and 75% of the IP samples showed no severe (PBC1 > 0.5) or moderate (PBC2 > 3) PCR bottlenecking level (**Fig. S1c and S1d**). Overall, all four indexes suggested that the library complexity of majority samples did not reach to a high level according to the ENCODE standards. Nevertheless, we note that some characteristics of RNA biogenesis are more complicated than DNAs, so new metrics may need to be developed for the evaluation of RNA library complexity.

Three peak calling tools --- exomePeak, MeTPeak and MACS2 --- have been widely used for m^6^A peak detection. We applied all three tools to identify m^6^A peaks from collected samples using fixed parameters. Jaccard Index (JI) and Simpson Index (SI) were adopted to assess the similarity of the peak sets identified by different tools. JI is defined as the number of intersecting bases between two peak sets divided by the number of bases in the union of the two peak sets[51] and SI measures the ratio of the number of intersecting bases between two peak sets to the number of bases in the smaller of the two peak sets[52]. Thus, by definition, for the same pair of peak sets, SI reaches a higher level due to its smaller denominator. To limit the comparisons at the transcriptome level, we only considered MACS2 peaks that were overlapped with annotated transcripts. Unexpectedly, only about 13.6% and 3.0% of the peak sets from MACS2 had 50% or larger complete overlap (JI > 0.5) with those from exomePeak and MeTPeak, respectively (**Fig. 3a and 3b**). This observation indicated poor reproducibility between peak sets called by MACS2 and those by exomePeak or MeTPeak. On the contrary, about 77.4% of the peak sets from exomePeak have JI > 0.5 with those from MeTPeak (**Fig. 3c**). In addition, about 73.0% and 86.6% of the peak sets from exomePeak have SI > 0.75 with those from MACS2 and MeTPeak, respectively (**Fig. 3a and 3c**). However, the proportion of the peak sets between MACS2 and MeTPeak with the same SI was reduced to 37.7% (**Fig. 3b**). It suggests that peaks called by MACS2 and MeTPeak achieve lower consistency than those by MACS2 and exomePeak. Taken together, exomePeak and MeTPeak agreed on over 75% of peak sets (JI > 0.5 or SI > 0.75), while MACS2 recovered limited peaks from them, especially from MeTPeak.

**Fig. 3.**
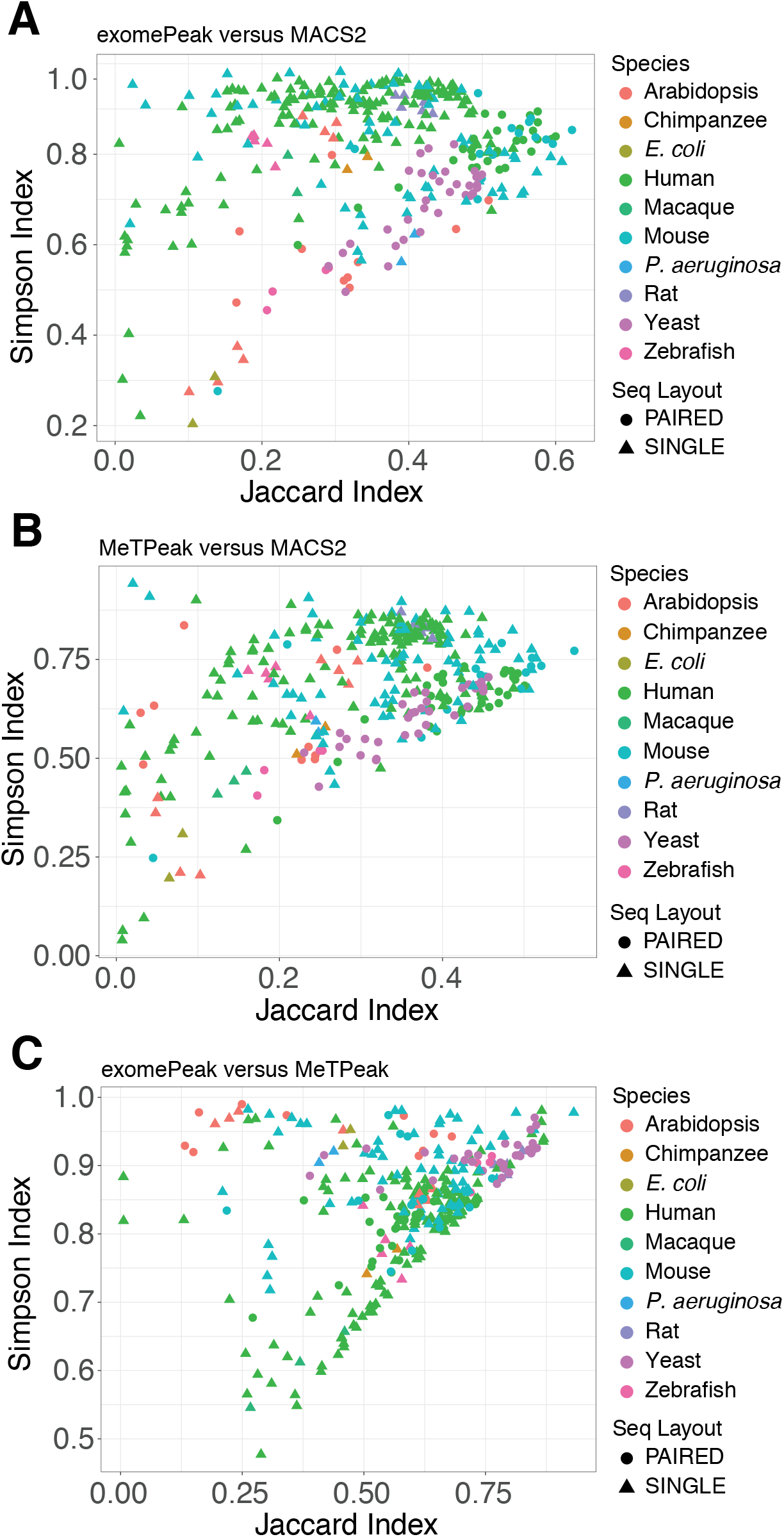
Evaluation of similarity of peak sets generated by three peak calling tools. Scatter plots showing the distributions of Jaccard index and Simpson index for comparisons of exomePeak versus MACS2 (a), MeTPeak versus MACS2 (b) and exomePeak versus MeTPeak (c) across all samples. Paired-end and single-end sequencing types are represented by triangle and circle, respectively. Species were indicated by colors.

### Cell- or tissue-specific m^6^A modifications

As genes are expressed in a tissue-specific manner, we ask whether m^6^A modifications possess similar characteristics. We first calculated the means of m^6^A peak fold enrichment levels in stop codon regions (±200 bp around the stop codons) of genes across cell lines and tissue types, and then examined top highest 2,000 coefficient of variations (CV) of these means. As a result, we observed strong correlations among samples from the same cell lines or tissue types (**Fig. 4a**), even they were collected from different studies/labs. Similarly, we observed in the t-distributed Stochastic Neighbor Embedding (tSNE)[53] plot that samples from the same cell or tissue type were clustered together and clearly separated from distinct types (**Fig. 4b**). These results suggest that some m^6^A modifications might regulate cellular activities and impact cellular processes in a cell line- or tissue type-specific manner, in response to different physiological conditions.

**Fig. 4.**
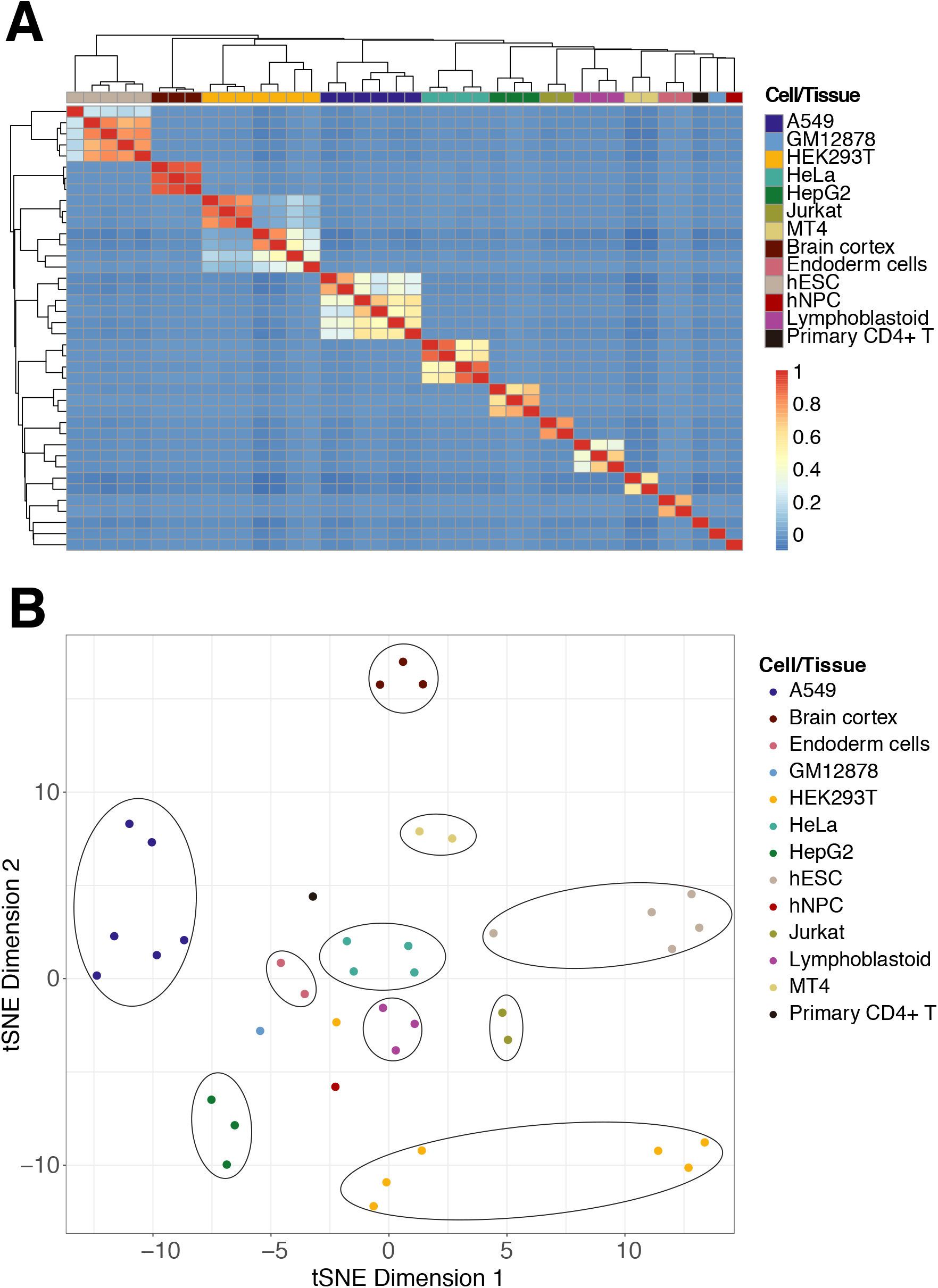
Cell- or tissue-specific m^6^A modifications. a) Heatmap plot depicting the Pearson correlation of different cell lines and tissues by using top highest 2,000 coefficient of variation (CV) of means of m^6^A peak fold enrichment levels at stop codon regions (±200 bp around the stop codons) of genes across cell lines and tissues. Cell/Tissue samples were clustered by complete linkage and the distances were measured by Euclidean distance. b) tSNE plot displaying clustering structure of different cell/tissue samples within the data used in (a). Each dot represents a sample and color indicates the cell/tissue type.

To offer insights into the cell line- or tissue-specificity of m^6^A modifications, REPIC (**Fig. 5a**) supports query of m^6^A modifications by cell lines or tissue types. On the Search page, we list options for all available cell lines and tissue types, next to filtering options including the number of peak sites in the gene of interest and samples from which peaks were called (**Fig. 5b**). Once the action of a query is complete, a report will be presented in a user-friendly interface with peak information including the genomic position, fold enrichment, genomic feature annotation (**Fig. S2a**) and sample information including the data source, read mapping statistics, metagene profiles and results from motif analysis (**Fig. S2b**).

**Fig. 5.**
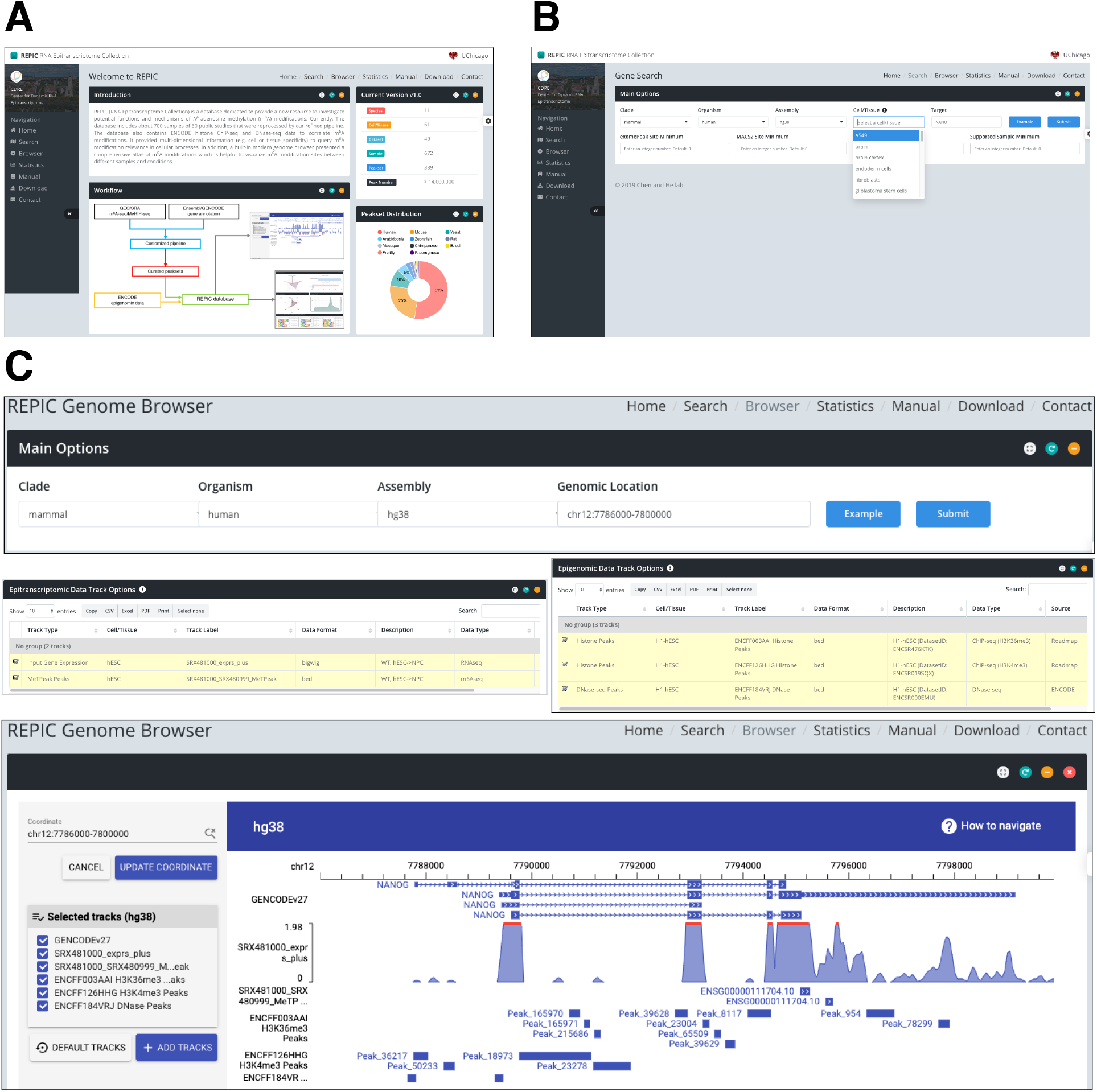
Screenshots of the web interfaces of the REPIC database. a) The home page. b) The search page. c) An example of visualizing m^6^A and histone modifications and chromatin accessibility in the gene NANOG. The top panel showed the query region including the upstream 2 kb of NANOG. The tracks for visualization were listed in the middle panel. The bottom panel displayed the result of the query in the genome browser.

### Visualization of m^6^A modifications and epigenomic data

The query on the Search Page is limited to genes. To better display multiple dimensional m^6^A modification information across the entire genome, REPIC provides a genome browser empowered by GIVE to visualize m^6^A peaks, fold enrichment and gene expression. As evidences unraveled that chromatin accessibility as well as epigenetic marks such as histone modifications define the cell/tissue types[54, 55], REPIC integrated DNase-seq and histone ChIP-seq data to investigate possible correlations between these epigenomic characteristics and m^6^A modifications. As a result, a total of 3,225 tracks including seven distinct track types (**Table S5**) constitute the built-in genome browser. Like the UCSC or other similar genome browsers, a user can select multiple tracks to interactively display peak or expression profile data at a specific genomic location. In an example demonstrating the utility of the browser shown in **Fig. 5c**, we observe H3K4me3 and DNase-seq peaks are located in the promoter region of the NANOG gene, indicating that it is actively transcribed in hESCs[12]. We also note that m^6^A modifications at the stop codon region are enriched with H3K36me3 peaks, which is consistent with the recently reported mechanism of m^6^A modification deposition[32].

### Future directions

As m^6^A modification detection technology has been applied to a variety of cell/tissue types with different conditions in distinct species, we will continue to collect new m^6^A/MeRIP-seq samples. In addition, with the increasing availability of transcriptome-wide sequencing data of m^6^A modifications at single-nucleotide resolution as well as other RNA modifications including m^1^A, m^5^C, m^7^G, Ψ and Nm, REPIC will be expanded to collect them. Another direction for the future development will be the integration of non-epitranscriptomic data such as RBP binding sites, GWAS, the GTEx data[56], etc to facilitate assessment and interpretation of RNA modifications.

## Conclusions

The current release of the REPIC database integrates millions of m^6^A peaks called by three popular tools from various cell/tissue types of multiple species. REPIC allows users to query m^6^A modification sites by specific cell lines or tissue types. Furthermore, hundreds of epigenomic datasets including chromatin accessibility and histone marks were added to the built-in genome browser to facilitate the interpretation of the functions of certain cell/tissue-specific m^6^A modifications, revealing their direct or indirect roles in influencing chromatin states and transcriptional regulation.

## Availability of data and materials

All 339 m^6^A peak sets called by our customized pipeline (https://github.com/shunliubio/easym6A) can be downloaded from the REPIC download center (http://epicmod.uchicago.edu/repic/download.php).

## Supplementary Table

**Table S1**. The list of 339 input-IP paired sample information. **Table S2**. The dataset list of histone ChIP-seq peaks from ENCODE. **Table S3**. The dataset list of DNase-seq peaks from ENCODE. **Table S4**. The genome assembly version and gene annotation source of 11 organisms. **Table S5**. The description of tracks in the genome browser.

## Supplementary Figure

**Fig. S1**. Library complexity of m^6^A-seq or MeRIP-seq data. **Fig. S2**. An example of the query of m^6^A modifications in a given gene.

