## Supplemental Figure for "REPIC: A database for exploring *N*^6^-methyladenosine methylome"

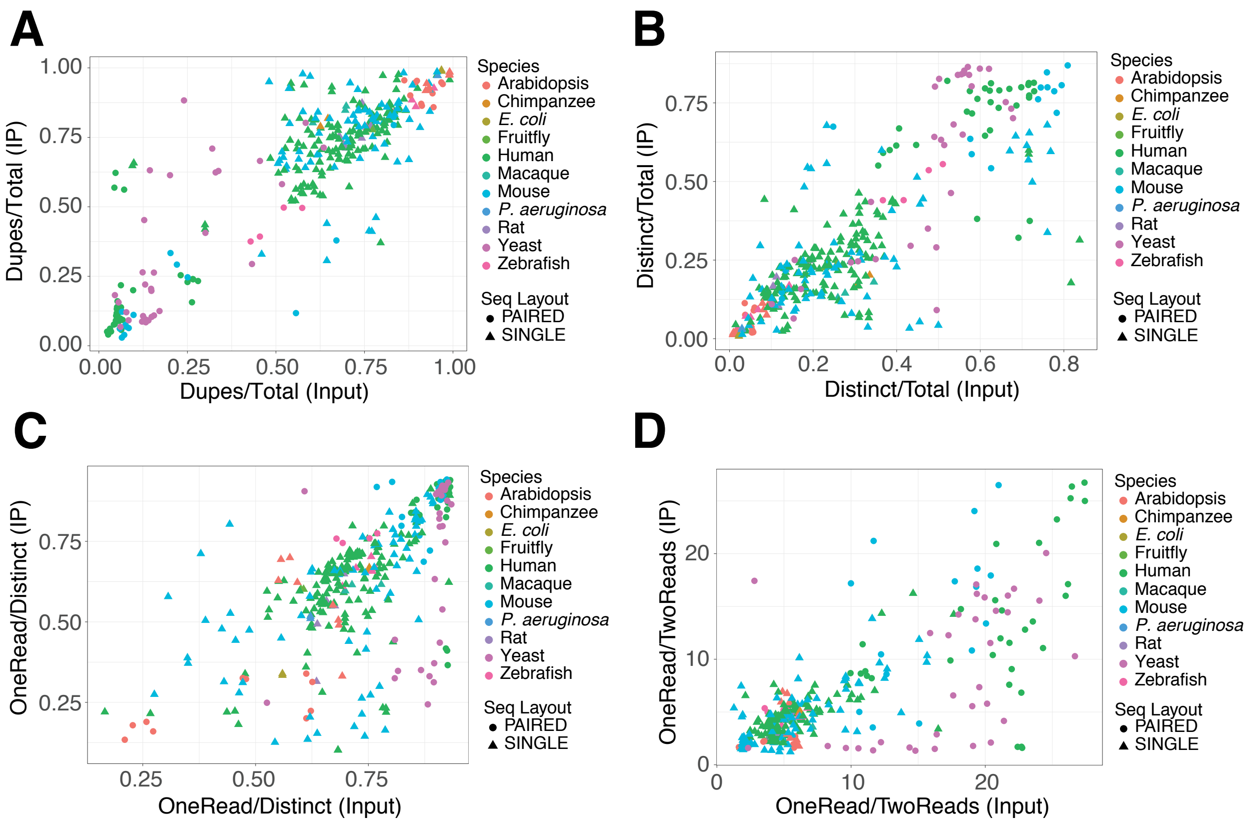


**Figure S1**. Library complexity of m$^6$A-seq or MeRIP-seq data. Scatter plots illustrating the distribution of input and IP samples within PDP (a), NRF (b), PBC1 (c) and PBC2 (d). Triangle and circle represent paired-end and single-end sequencing data type, respectively. Color indicates species.

A

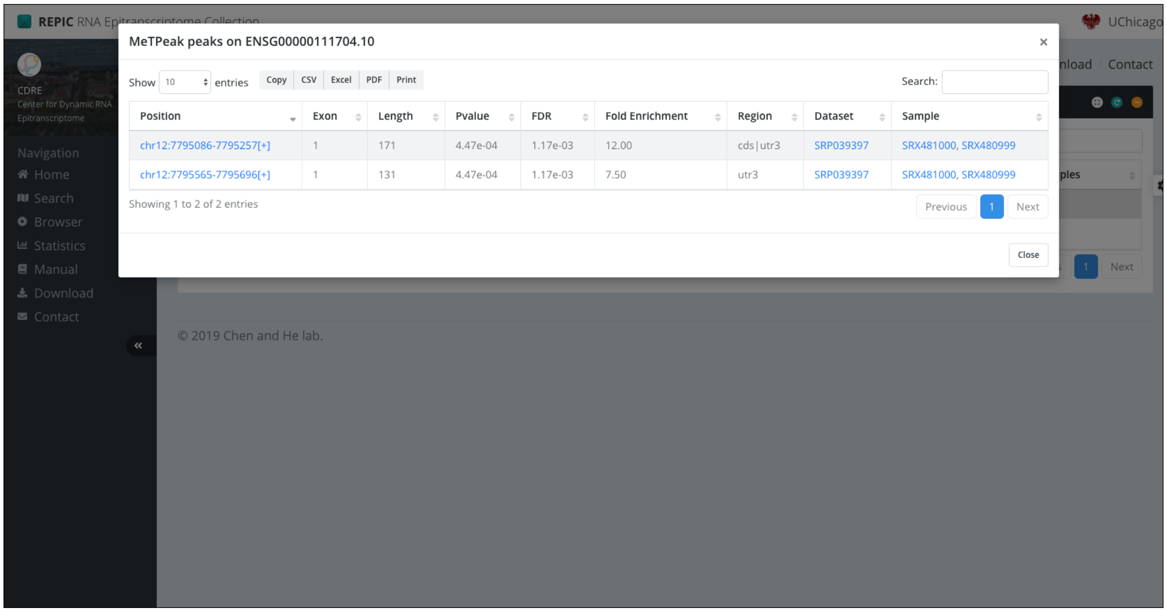

REPIC RNA Epitranscriptome Collection
MeTPeak peaks on ENSG00000111704.10
Show 10 entries
Copy
CSV
Excel
PDF
Print
Search:
Position
Exon
Length
Pvalue
FDR
Fold Enrichment
Region
Dataset
Sample
chr12:7795086-7795257[+]
1
171
4.47e-04
1.17e-03
12.00
cds|utr3
SRP039397
SRX481000, SRX480999
chr12:7795565-7795696[+]
1
131
4.47e-04
1.17e-03
7.50
utr3
SRP039397
SRX481000, SRX480999
Showing 1 to 2 of 2 entries
Previous
Next
Close
CDRE
Center for Dynamic RNA Epitranscriptome
Navigation
Home
Search
Browser
Statistics
Manual
Download
Contact
UChicago
© 2019 Chen and He lab.


B

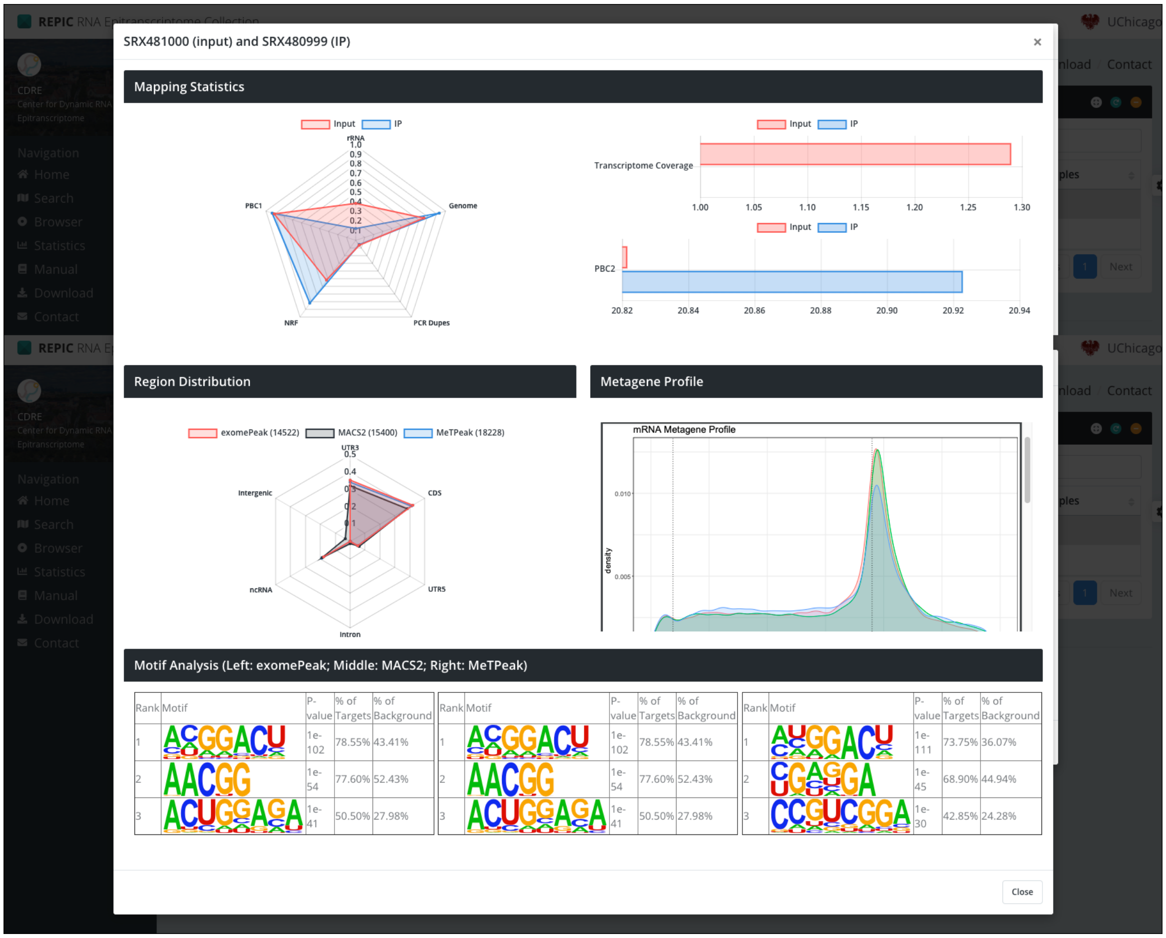

REPIC RNA Epitranscriptome Collection
SRX481000 (input) and SRX480999 (IP)
Mapping Statistics
Input
IP
rRNA
Genome
PCR Dupes
NRF
PBC1
Transcriptome Coverage
PBC2
Region Distribution
exomePeak (14522)
MACS2 (15400)
MeTPeak (18228)
UTR3
CDS
UTR5
Intron
ncRNA
Intergenic
Metagene Profile
mRNA Metagene Profile
density
Motif Analysis (Left: exomePeak; Middle: MACS2; Right: MeTPeak)
Rank
Motif
P-value
% of Targets
% of Background
1e-102 78.55% 43.41%
1e-54 77.60% 52.43%
1e-41 50.50% 27.98%
1e-111 73.75% 36.07%
1e-45 68.90% 44.94%
1e-30 42.85% 24.28%
Close

**Figure S2**. A query of $m^6A$ modifications in a given gene. Screenshots of the web interfaces for a practical use to query $m^6A$ modifications in an interested gene (e.g. NANOG) from different cell lines or tissues (a) and to browse information generated by data processing in associated samples (b).
